# Supervised Deep Learning with Gene Annotation for Cell Classification

**DOI:** 10.1101/2024.07.15.603527

**Authors:** Zhexiao Lin, Yuanyuan Gao, Wei Sun

## Abstract

Gene-by-gene differential expression analysis is a widely used supervised approach for interpreting single-cell RNA-sequencing (scRNA-seq) data. However, modern scRNA-seq datasets often contain large numbers of cells, which can produce numerous differentially expressed genes with exceedingly small p-values but minimal effect sizes, and thus making biological interpretation difficult. To overcome this challenge, we developed Supervised Deep learning with gene ANnotation (SDAN), a method that integrates gene-annotation information with gene-expression profiles using a graph neural network. SDAN identifies functionally coherent gene sets that best classify cells, and the resulting cell-level classification scores can be aggregated to make individual-level predictions. We evaluated SDAN and two representative existing methods in three real-data applications to identify gene sets associated with severe COVID-19, dementia, and immunotherapy response in cancer. SDAN consistently outperformed alternative approaches by achieving two key objectives simultaneously: accurate classification of outcomes and unambiguous assignment of genes to gene sets of functionally related genes.

## Introduction

A key step in analyzing single-cell RNA-seq (scRNA-seq) data is gene-by-gene differential expression (DE) analysis, often followed by identifying biological processes enriched among the DE genes. However, due to the large number of cells in modern scRNA-seq datasets, DE analyses frequently yield extremely small p-values for a substantial number of genes, even when the corresponding effect sizes are minimal. As a result, researchers must apply additional ad hoc filtering to obtain interpretable results. To address this challenge, we focus on directly identifying gene sets that accurately classify cells. Gene sets offer more interpretable biological insights than long lists of DE genes, and classification accuracy is often more meaningful for practical applications than p-values.

To achieve this goal, we developed Supervised Deep learning with gene ANnotation (SDAN), a graph neural network (GNN)–based method that integrates gene-annotation information with gene-expression data. SDAN builds a gene–gene interaction graph to represent annotation relationships, incorporates gene expression as node attributes, and learns gene sets whose averaged expression profiles classify cells and whose member genes show coherent expression patterns and proximity within the graph. Unlike conventional neural network models that operate as “black boxes”, SDAN provides inherently interpretable outputs in the form of gene sets, enabling transparent and biologically meaningful interpretation of the learned representations.

A central strength of deep learning lies in representation learning, the ability to derive new features that are often complex functions of the original inputs. In scRNA-seq analysis, however, the original features (genes) are already biologically meaningful and inherently interpretable, limiting the utility of replacing them with latent representations. This may help explain why most deep learning applications in scRNA-seq have centered on unsupervised tasks, including dimension reduction and denoising [1–3], clustering [4, 5], and batch correction or harmonization [6, 7], rather than on supervised prediction. SDAN can be considered as a combination of un-supervised clustering of genes and supervised classification of cells. The model learns gene clusters that are directly optimized to improve classification performance while ensuring that the derived gene sets are both interpretable and task-relevant.

Several factorization-based methods aim to identify gene sets that capture shared patterns in single-cell data. Although these approaches do not incorporate supervised classification, the gene sets they produce can be used to train downstream classifiers, allowing a fair comparison with SDAN. Broadly, factorization methods can be grouped by whether they utilize gene annotations. Among those that do not use gene annotation, a representative method is consensus non-negative matrix factorization (cNMF) [8]. However, CMF is computationally expensive, computationally efficient method for matrix factorization, such as sciRED [9] has been developed for large-scale scRNA-seq datasets. The second group of factorization methods incorporate gene annotation. A representative example is Spectra [10], which models both cell type–specific and cell type–agnostic factors. We evaluated SDAN, sciRED, and Spectra across three real-world classification tasks: distinguishing severe from mild COVID-19, dementia from healthy controls, and cancer immunotherapy responders from non-responders. In all cases, the ultimate objective is to classify individuals, although predictions are first made at the cell level and then aggregated to produce an individual-level label.

## Results

### Overview of SDAN

SDAN can be considered as a two-step method (Figure 1). In the first step, we combine scRNA-seq data and gene annotations to identify latent components in scRNA-seq data using a GNN. Each component is a linear combination of all the genes. The latent components are estimated through a graph pooling operation of the GNN, where nodes (genes) with similar expression and similar annotations (i.e., connected in the graph) are pooled into one latent component. To facilitate interpretation, a penalty term is included in the objective function to encourage sparse assignments, making the assignments of genes to latent components almost binary. Therefore, these latent components can be considered as gene sets. In the second step, we project scRNA-seq data to these gene sets and then use the gene-set-level data to classify cells by a multilayer perceptron (MLP). These two neural networks (GNN for latent component identification and MLP for outcome prediction) are trained together.

**Figure 1:**
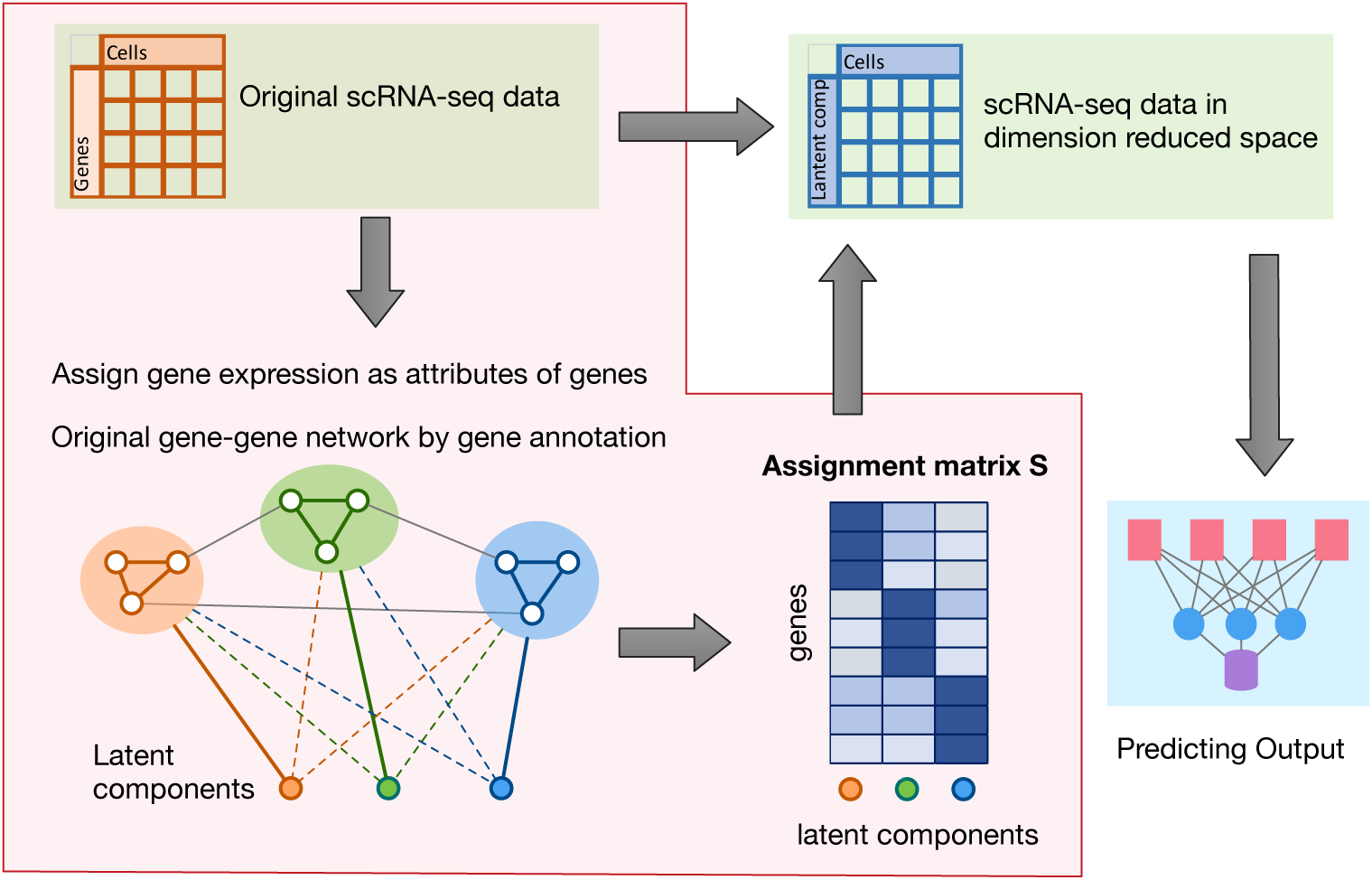
An overview of Supervised Deep Learning with Gene Annotation (SDAN) can be divided into two steps. In the first step (shaded area), SDAN combines gene expression data and gene annotations (gene-gene interaction graph) to learn a gene assignment matrix **S**, which specifies the weights of each gene for all latent components. In the second step, SDAN uses the gene assignment matrix **S** to reduce the gene expression data of each cell to a low-dimensional space (from the number of genes to the number of latent components) and then makes predictions in the low-dimensional space using a feed-forward neural network.

The loss function of SDAN includes an unsupervised component and a supervised component. The unsupervised loss function is the loss for the GNN, which encourages genes connected in the graph to be included in the same gene set. The supervised loss function is the cross-entropy loss for classification accuracy. See the Online Methods Section for details. We adjusted the contributions of the two losses by adding a weight for the unsupervised loss and explored a series of values for this weight from 0 to 10. Note that when this weight is 0, it becomes a baseline approach without using gene annotations. We provided guidance on how to choose the weight and we found that weight 2 was a robust and optimal choice in all the datasets that we analyzed, since it resulted in accurate predictions and strong functional coherence among genes within each gene set.

### Gene expression of CD8+/CD4+ T cells can classify severe versus mild COVID-19 patients

Several studies have shown that T cell response induced by vaccination or natural infection is crucial in protecting against severe COVID-19 [11–13]. An ineffective or delayed T cell response may lead to over-activation of the innate immune system, causing tissue damage. In contrast, a timely and effective T cell response can clear the virus and thus prevent severe COVID-19. We conjecture that T cells that react to SARS-CoV-2 have gene expression patterns that can distinguish them from other T cells. We used scRNA-seq data from ∼ 42,000 CD4+ T cells and ∼ 24,000 CD8+ T cells from 49 mild and 32 severe COVID-19 patients [14], and split the data so that half of the patients were used for training and the other half for testing. Furthermore, 10% of the cells in the training data were used as validation data to decide when to stop deep learning training. See Supplementary Materials for more details.

Next, we illustrated the consequence of different weights on unsupervised loss while predicting severe COVID-19 by CD8+ T cell gene expression. The conclusion was similar when we explored other applications. SDAN estimates 40 latent components (gene sets) by default. When the weight for the unsupervised loss is small, many components are empty, i.e., without contributing genes. As the weight for the unsupervised loss increases, the number of non-empty latent components increases and stabilizes at 40 (Figure 2(A)). We assessed whether the genes within a latent component were more connected than expected by chance using a quantile measurement. Denote the connectivity of an observed gene set *G* by *C_G_*, which is the average degree in the gene set, i.e., 2 × number of edges/number of genes. We randomly selected many gene sets of the same size and calculated the distribution of their connectivities, denoted by *f* (*G*). Then we calculated what quantile of *f* (*G*) is *C_G_*. As expected, as the weight for the unsupervised loss increases, the quantile become larger (Figure 2(B)). Finally, we aggregated the cell-level prediction scores to individual-level prediction scores by taking the average, and used the individual level prediction scores to predict COVID-19 severeness. The accuracy of the prediction decreases slightly as the weights for the unsupervised loss increase (Figure 2(C)). The CD4+ T and CD8+ T celllevel AUCs vary from 0.89 to 0.85 and from 0.91 to 0.83 respectively, though individual level AUC are more stable and are between 0.94 and 0.97 for both cell types (see Supplementary Materials Section 1.2 for more details).

**Figure 2:**
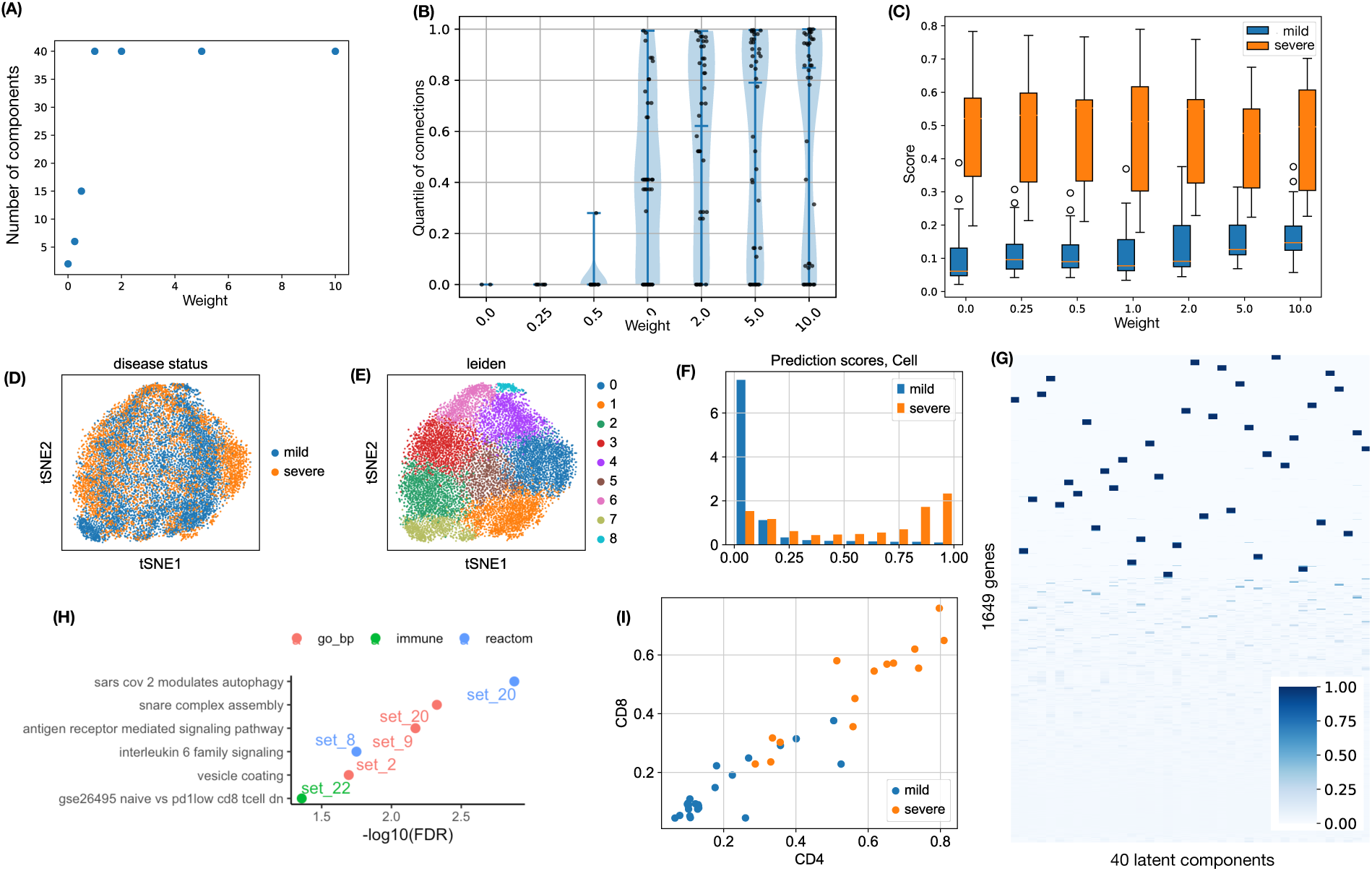
Summary of results for CD8+ T cells. **(A)** The number of latent components for different weights for the unsupervised loss. **(B)** Violin plots of connection quantiles. Each point represents one component. The connection quantile is computed based on the number of connections from randomly selected gene sets. **(C)** Boxplots of individual-level scores (average of cell level scores) for severe disease, across different weights for the unsupervised loss. **(D-E)** tSNE projection of CD8+ T cells in testing data, colored by disease status or clusters generated by the Leiden algorithm. **(F)** Distribution of cell-level prediction scores (for CD8+ T cells using weight 2.0) for severe disease in testing data, stratified by the status of severe/mild diseases of the corresponding individual. **(G)** Assignment matrix of 1,649 genes to 40 latent components. Genes are ordered by hierarchical clustering. **(H)** Enriched functional categories of identified gene sets. **(I)** Individual level scores COVID-19 patients calculated based on gene expression of CD4+ or CD8+ T cells.

While balancing the prediction accuracy and the utilization of gene annotation, we chose to use a weight 2 for the unsupervised loss. Clustering analysis of CD8+ T cells cannot reveal disease status (Figure 2 (D-E)). In contrast, the SDAN prediction scores are very different between the cells from severe patients and the cells from mild patients (Figure 2 (F)). Further analysis shows several components are enriched with functional categories from gene ontology (biological processes), reactome pathways, or immunologic gene sets (Figure 2 (G-H)). For example, gene set 20 is enriched with genes from the reactome pathway of SARS-Cov-2 modulates autophagy, and GO term SNARE complex assembly. Autophagy is activated during SARS-Cov-2 infection to destroy the infected cells. However, some proteins encoded by SARS-Cov-2 can prevent this process by blocking the assembly of the SNARE complex, which is required for the autophagy process [15]. The enriched functional categories are similar for larger weights for the unsupervised loss (Supplementary Figure 2).

The results from CD4+ T cells also support the use of weight 2 for the unsupervised loss (Supplementary Figure 3). Individual-level prediction scores made by CD8+ T cells and CD4+ T cells are highly consistent (Figure 2 (I)).

We have used the average of cell-level prediction scores to calculate individual-level prediction scores. While this is a reasonable approach, it is important to explore the variation of cell level prediction scores. When using either CD8+ T cells (Figure 2(F)) or CD4+ T cells (Supplementary Figure 3(D)) to predict severe COVID-19, the T cells from mild patients tend to have smaller prediction scores concentrating around 0, and the prediction scores from severe patients have a peak around 1, but also spread in the range from 0 to 1. This implies that a subset of T cells from severe patients might be responsible for disease progression.

### Gene expression of astrocytes or microglia can classify dementia status

Alzheimer’s disease, which is associated with the accumulation of amyloid beta plaques and hyperphosphorylated tau, is the most common cause of dementia in older individuals [16]. Many recent studies have pointed out that the human immune response plays an important role in the initiation, progression, and pathology of Alzheimer’s. Two types of non-neuronal cells, microglia and astrocyte, are involved in this process [17, 18]. We used SDAN to analyze the snRNA-seq data from the Seattle Alzheimer’s Disease Cell Atlas (SEA-AD) consortium [19], focusing on ∼ 70,000 astrocyte (Astro) cells and ∼ 40,000 microglia/perivascular macrophage (micro-PVM) cells from 84 donors (42 with dementia and 42 without dementia). We sought to classify dementia status using astrocyte or micro-PVM expression. Similar to the COVID-19 study, we randomly selected half of the individuals as training data and used the remaining ones as testing data. We also evaluated the results across different weights for the unsupervised loss and concluded that a weight of 2.0 for the unsupervised loss is an appropriate choice (Supplementary Figures 3-4). We further explored in detail the results with a weight of 2.0.

In contrast to predicting severe COVID-19 using T cells, predicting dementia using astrocyte or microglia-PVM is more challenging. This can be illustrated by comparing cell level and gene level prediction scores in Figure 2(C) and (F) (CD8+ T cells to predict COVID-19) and Figure 3(A-D). Many astrocyte and microglia-PVM cells have similar prediction scores between dementia and non-dementia donors (Figure 2(A-B)), though a subset of them have higher scores in dementia donors. The individual-level prediction scores cannot separate dementia and non-dementia donors very well (with AUC 0.744 and 0.735 for astrocyte and microglia-PVM, respectively; see Supplementary Materials Section 2 for more details). However, when jointly considering the prediction scores by astrocyte and microglia-PVM, a subset of donors stands out. They have high scores using either astrocyte or microglia-PVM (Figure 3(E)). We refer to this subset of dementia donors as dementia-i where i stands for immune system, and refer to the remaining dementia donors as dementia-s, where s stands for immune <u>s</u>ilence.

**Figure 3:**
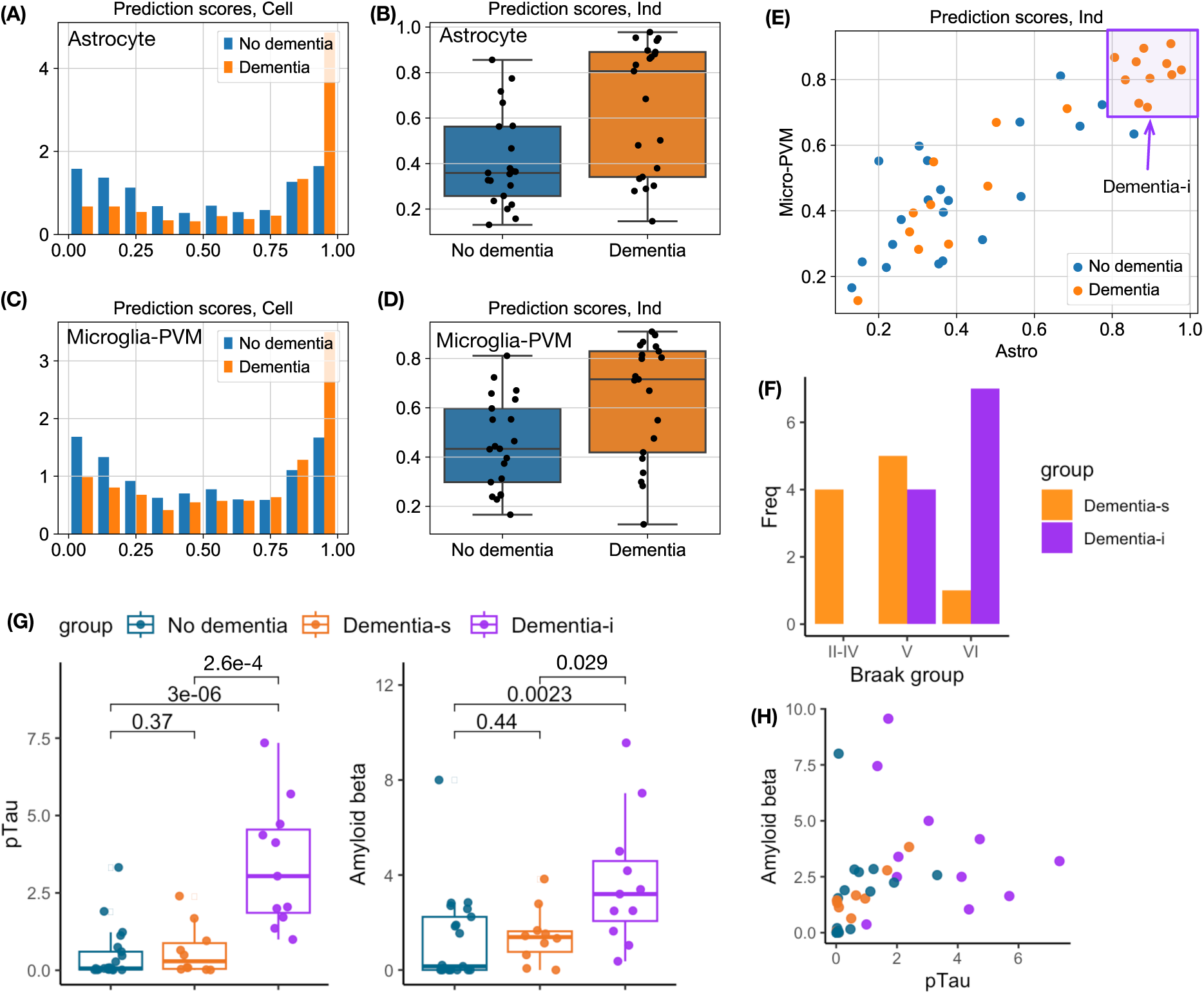
**(A-B)** The distributions of cell level and individual level prediction scores for dementia using astrocyte in testing data. **(C-D)** The distributions of cell level and individual level prediction scores for dementia using microglia-PVM in testing data. **(E)** Scatter plot of individual level scores by astrocyte or microglia-PVM, in testing data. We identified a Dementia subset. **(F)** The Braak staging groups for donors who belong to the Dementia subset or the remaining Dementia donors. **(G)** Percentage of pTau (Amyloid beta) positive area for three groups of donors: No dementia, Dementia-s, and Dementia-i. **(H)** Scatter plot of the percentage of pTau positive area vs. the percentage of Amyloid beta positive area.

We further characterize the dementia-i donors by the neuropathology measurements. Hyperphosphorylated tau aggregates into insoluble twisted fibers known as neurofibrillary tangles. Braak staging is a popular system used to classify the extent of neurofibrillary tangle pathology in Alzheimer’s disease. The staging system is divided into six stages with stage IV being the most severe. All four donors at Braak stage II-IV are dementia-s; most (7 out of 8) donors at Braak stage VI are dementia-i, while donors at Braak stage V are a mixture of dementia-s and dementia-i (Figure 3(F)). These results suggest that dementia-i donors have more advanced diseases.

This set of dementia-i donors also have much higher pTau and amyloid beta measurements than dementia-s or non-dementia donors (Figure 3(G)), while pTau and amyloid beta were measured as the percentage of areas that are positive for pTau (phosphorylated tau, measured by antibody AT8) or Amyloid beta (by antibody 6e10) by Gabitto et al. [19]. Combining pTau and amyloid beta measurement may give a better characterization of the dementia-i donors (Figure 3(H)), though a larger sample size is needed to confirm this conclusion.

### Gene expression of CD8+ T cells can predict cancer patients’ response to immunotherapy

Cancer immunotherapy, particularly immune checkpoint inhibitor (ICI), has revolutionized cancer treatment, yet predicting which patients will respond to it remains a significant challenge [20]. The goal of ICI is to induce or strengthen the T cell response to tumors. Previous studies have demonstrated that CD8+ T cell gene expression is informative in predicting patient responses to ICI [21, 22]. To address this critical and challenging problem, we employed SDAN to identify gene sets associated with ICI response. Due to the limited sample sizes in most existing studies, we used scRNA-seq data from one study, consisting of 6,350 CD8+ T cells from 17 responding tumors and 31 non-responding tumors [21], as our training data. For testing, we used scRNA-seq data from Yost et al. [22], which included 27,924 CD8+ T cells from 8 responders and 7 non-responders.

Direct clustering of CD8+ T cells cannot separate cancer patients who respond to ICI from those who do not (Figure 4 (A)). In the testing data by Yost et al. [22], the classification accuracy at the cell level is low, with AUC in the range of 0.52 to 0.59 for unsupervised loss weights from 0 to 10. After aggregating information across cells, the individual-level AUC varies from 0.48 to 0.82 (Supplementary Materials Section 3). A weight of 2.0 for the unsupervised loss is still a good choice (Supplementary Figure 5). At this weight, the cell-level and individual-level AUCs are 0.53 and 0.66, respectively (Figure 3B).

**Figure 4:**
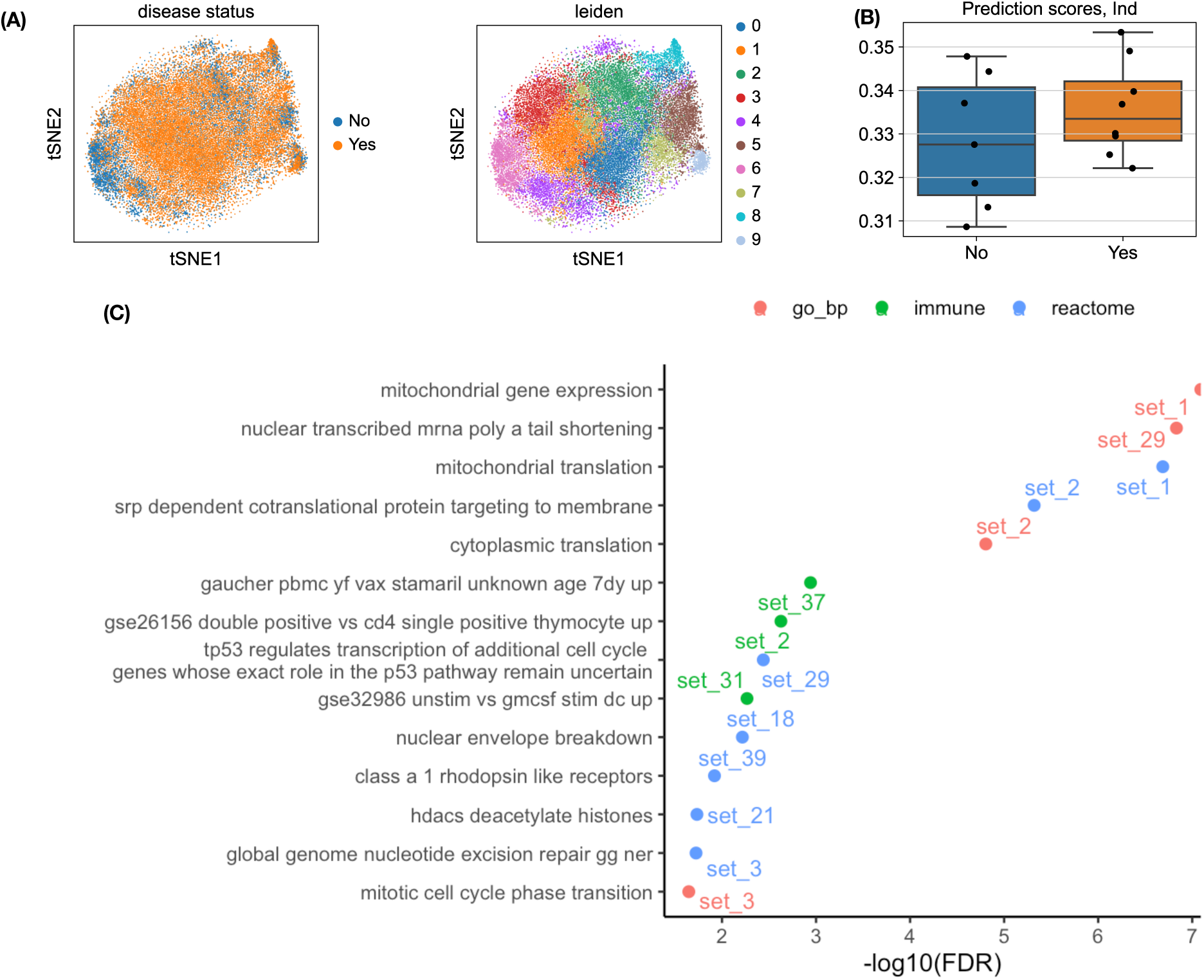
**(A-B)** tSNE projection of CD8+ T cells in testing data, colored by the response to immunotherapy (Yes or No) or clusters generated by the Leiden algorithm. **(B)** The distributions of individual level prediction scores for patients who respond to immunotherapy (Yes) or not (No), using weight 2.0 for unsupervised loss. **(C)** The functional categories that are over-represented by the genes in the 40 gene sets learned by SDAN, using a weight of 2.0 for unsupervised loss.

Next, we queried the functional categories that were enriched in the gene sets identified by SDAN. The top enrichment is from gene set 1, which is enriched with genes involved mitochondrial function. This enrichment is robust for unsupervised weights from 2.0 to 10 (Supplementary Figures 6-7). Gene set 1 is composed of 23 genes, including six genes that are mitochondrial ribosomal proteins or involved in mitochondrial ribosome function, three genes involved in transcription/translation in mitochondrial, a mitochondrial methionyl-tRNA formyltransferase (MTFMT), two mitochondrial tRNA synthetases ß (SARS2 and EARS2), and four genes involved in NADH:ubiquinone oxidoreductase (Supplementary Table 1). NADH plays an essential role in generating energy within cells. The collection of mitochondrial and NADH genes suggests that this gene set captures the metabolism characteristics of T cells. It is well recognized that the metabolism state of T cells is closely related to T cell immune response [23]. Recent studies have also shown that mitochondrial dysfunction could affect T cell response to neoantigens (e.g., leading to terminally exhausted T cells) [24, 25], and thus may affect cancer patients’ response to immunotherapy.

### Comparison of SDAN vs. Spectra and sciRED

We compared the performance of SDAN with two representative existing methods, Spectra [10] and sciRED [9]. Because neither Spectra nor sciRED is explicitly designed for supervised learning, we conducted a fair comparison by evaluating the gene sets identified by all three methods within the same downstream framework. Specifically, for each method, we applied factor analysis to the gene expression data to obtain latent components, fit a logistic regression model for cell-level classification using the resulting components, and then aggregated the cell-level predictions to perform individual-level classification. See Methods Section for implementation details.

All three methods achieve comparable performance in cell-level classification as well as in individual-level classification based on aggregated cell-level prediction scores (Supplementary Tables 2–5). However, the gene sets identified by the three methods differ substantially in their structure and interpretability. In particular, sciRED is designed to learn latent components without enforcing sparsity on gene weights. As a result, each latent component typically involves thousands of genes with nonzero weights. This lack of sparsity makes the resulting gene sets difficult to interpret and less suitable for downstream biological analyses (Figure 5, Supplementary Figure 8).

**Figure 5:**
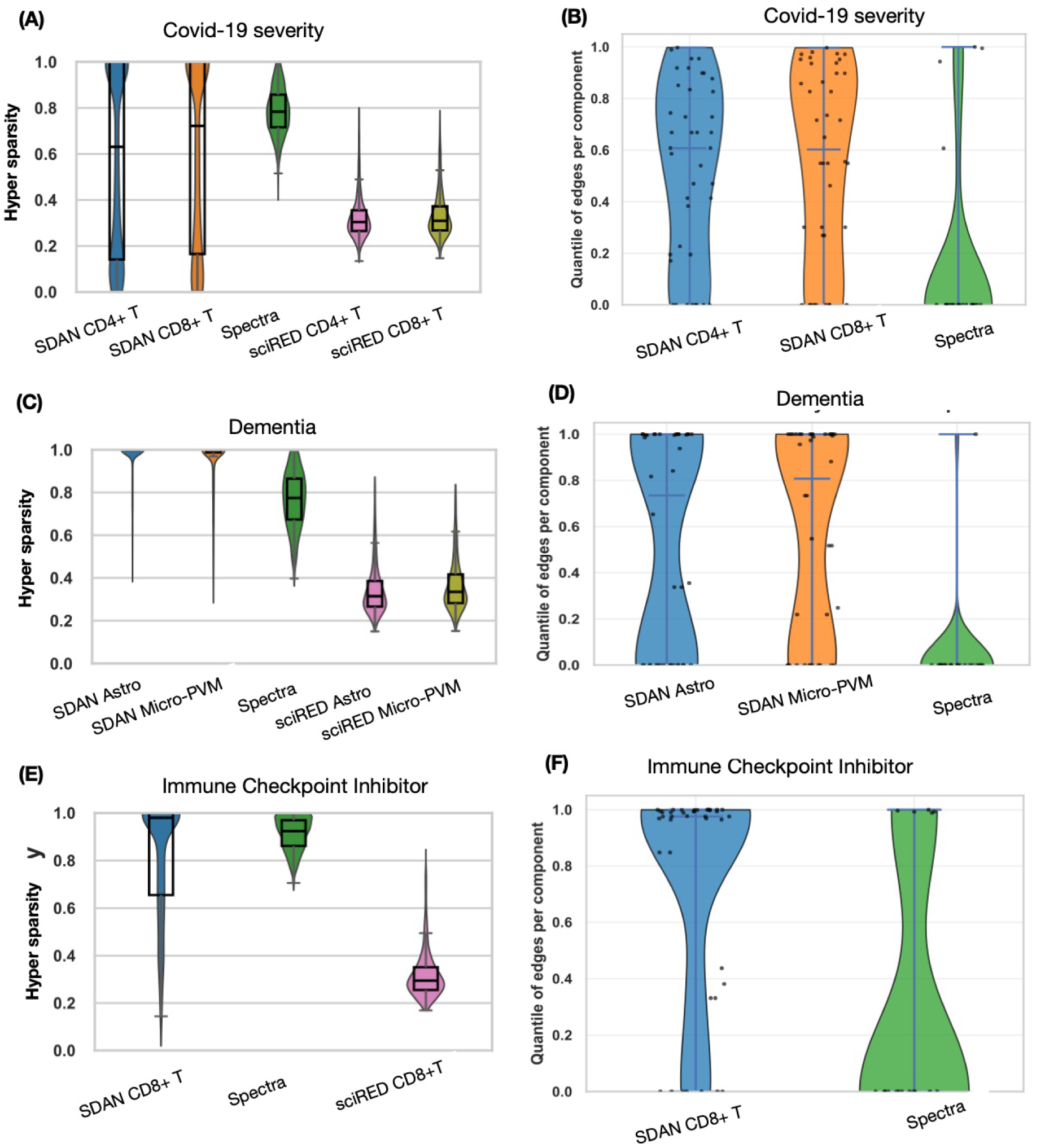
**(A, C, E)** Comparison of sparsity of gene loading vectors. We quantify the sparsity of the loading vector for each latent component using Hoyer sparsity. Hoyer sparsity takes a value of 0 if all entries are equal (maximally dense), and a value of 1 if there is only one nonzero entry (maximally sparse) (See Supplementary Section 4 for details). The violin plots and box-plots summarize the Hoyer sparsity of all latent components identified by each method. Because Spectra internally accounts for cell-type specificity, its components are not cell-type specific. **(B, D, F)** Connectivity enrichment of inferred gene sets. We assess whether genes within each inferred gene set exhibit more connections in the gene–gene interaction knowledge graph than expected by chance. A reference distribution is generated by repeatedly sampling random gene sets of the same size, and the connectivity enrichment of each component is quantified by its quantile within this reference distribution.

Although the latent components identified by Spectra enforce sparsity on gene weights and therefore involve substantially fewer genes than those identified by sciRED, they remain less well defined than the latent components inferred by SDAN (Figure 5, Supplementary Figures 9–10). Because both SDAN and Spectra leverage the same gene–gene interaction annotations to construct gene sets, we directly compared the within–gene-set connectivity under the annotation graph for the two methods. On average, gene sets identified by SDAN exhibit higher connectivity in the underlying annotations than those identified by Spectra, indicating that SDAN more effectively recovers functionally coherent gene modules.

## Discussion

Biological knowledge, such as gene-gene interactions, is often noisy and may vary across different tissues or conditions. Consequently, when using such knowledge in machine learning methods, it is crucial to have the flexibility to learn the subset of biological knowledge that is relevant to the particular task. The SDAN method achieves this goal by combining supervised classification loss and unsupervised graph loss, which prefer gene sets that align with biological knowledge and their expression can classify cells. In contrast, many earlier works incorporate biological knowledge into neural networks by designing the neural network structure using gene-regulation networks [26–29] and lack the flexibility to select the relevant subset of biological knowledge.

In this work, we label cells according to their corresponding individuals. For example, when comparing cells from severe and mild COVID-19 patients, we label the cells from individuals with severe or mild cases as severe or mild cells, respectively. However, some of these cell labels, inherited from their corresponding individuals, may be incorrect due to cellular heterogeneity. Despite these noisy labels, we have demonstrated that SDAN can still learn and extract useful signals. An interesting future direction is to explicitly handle such noisy labels [30–32].

We compared SDAN with two alternative methods, sciRED [9] and Spectra [10]. In cell-level differential expression and classification analyses, high classification accuracy is generally achievable, in part due to the large number of cells available. Consistent with this expectation, all three methods achieve similar classification accuracy across all three datasets. However, SDAN outperforms the other two methods in terms of interpretability by identifying well-defined gene sets that are enriched with internal connectivity in known gene–gene annotation networks.

## Methods

### Neural network architecture and loss functions

Let *n* be the number of cells in the training data, *p* be the number of genes, and *d* be the number of components after dimension reduction. Each gene is regarded as a node in the gene-gene interaction graph, with the node features being the expression data for each gene. We utilized a graph pooling operator to reduce the gene expression from *p* genes to *d* components. Denote the transposed gene expression data matrix by *X* ∈ ℝ*^p^*^×*n*^. Gene annotations are summarized by an adjacency matrix *A* ∈ ℝ*^p^*^×*p*^ (including self-connections). Some annotations, such as protein-protein interactions, are already in this adjacent matrix form. For other annotations that group genes into categories, such as gene ontology, genegene similarities may also be defined as a continuous measurement [33], and thus create a weighted gene-gene interaction graph. For our method, *A* does not need to be binary; it can also represent a continuous measurement of similarity. Let *Ã* be the normalized adjacency matrix *Ã* = *D*^−1*/*2^*AD*^−1*/*2^, where *D* is the degree matrix of *A*. The goal is to learn an assignment matrix *S* ∈ ℝ*^p^*^×*d*^ to assign some of the *p* genes to the *d* components.

The GNN has two steps: graph convolution followed by graph pooling. We apply the graph convolution networks [34] to reduce the high-dimensional data of each gene (i.e., its expression in *n* cells) to a small number of hidden states *h*:

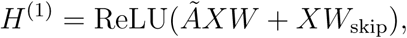

where *H*^(1)^ ∈ ℝ*^p^*^×*h*^ is the output of the first layer, *W* ∈ ℝ*^n^*^×*h*^ and *W*_skip_ ∈ ℝ*^n^*^×*h*^ are two weight matrices to be estimated, and ReLU denotes the non-linear activation function Rectified Linear Unit. *ÃX* represents the graph-convoluted gene expression data. The same weight matrix *W* is applied to all the genes, resembling the operation in a convolutional neural network where the same convolutional operation is applied across all segments of an image. The additional *XW*_skip_ allows each gene to give more weight to its own expression. This graph convolution layer can be applied multiple times. For the *ℓ*-th layer, the formulation is *H*^(*ℓ*)^ = ReLU(*ÃH*^(*ℓ*−1)^*W* ^(*ℓ*−1)^ + *H*^(*ℓ*−1)^*W*_skip_^(*ℓ*−1)^). The output of the final layer is denoted by *H*. In our implementation, we used two hidden layers for the graph convolution networks, with the number of hidden states being 64 in each layer.

The second step is graph pooling. The simplest case is one-step graph pooling. The assignment matrix *S* is obtained by *S* = Softmax(*HW_H_*) ∈ ℝ*^p^*^×*d*^, where the Softmax operation is applied on each row (i.e., each gene) to assign the weights of each gene to *d* latent components, and *W_H_* ∈ ℝ*^h^*^×*d*^ is a weight matrix to be estimated. Graph pooling can also involve multiple layers, which becomes hierarchical graph pooling. Let *X_r_* = *X*^⊤^*S* be the projection of scRNA-seq data in the latent space. We used *X_r_* to predict class labels via a MLP with three hidden layers, each containing 64 hidden nodes and a ReLU activation function.

Now we define the loss function as a combination of two components: an unsupervised graph loss and a supervised classification loss. For the graph loss, we adopt the “minCUT” loss, which is formulated to resemble spectral clustering [35]. The graph loss is the sum of two terms: ℒ_c_ and ℒ_o_, defined as follows:

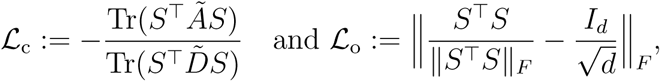

where ‖·‖*_F_* denotes the Frobenius norm, *Ã* is the normalized adjacency matrix, *D̃* is the degree matrix of *Ã*, and *I_d_* is the *d*-dimensional identity matrix. The term ℒ_c_ encourages strongly connected nodes to be grouped together. The term ℒ_o_ encourages gene assignment to be orthogonal (i.e., each gene is assigned to only one latent component) and ensures that all latent components have similar sizes. The classification loss is defined by the crossentropy loss, denoted as ℒ_clf_. Therefore, the final loss function is ℒ_clf_ + *w*(ℒ_mc_ +ℒ_o_), where *w* adjusts the weight for the unsupervised graph loss. We set *w* = 2 by default.

### Evaluation and generalization

Let *X* ∈ ℝ*^n′^*^×^*^p′^* be the test data, where *p*^′^ may not be equal to *p*; that is, the gene lists of the training data and the test data may be different. When the gene list of the test data is the same as that of the training data, the dimension reduction of the test data is performed as *X*_test,r_ = *X*_test_*S* using the trained assignment matrix *S*. We then apply the trained MLP to *X*_test,r_ to obtain the scores for each cell in the test data. If the gene list of the test data differs from that of the training data, we identify the common genes between the test data and the training data. We then use the corresponding sub-matrices of *X*_test_ and *S* to perform the dimension reduction on test data.

### Implementation of SDAN

The gene-gene interaction graph was constructed based on protein-protein interaction annotation from the BioGRID database: `BIOGRID-ORGANISM-Homo_sapiens-4.4.204.tab3.txt.gz`. We matched genes by their official symbols in the BioGRID database. There is an edge between two genes if their interaction exists in BioGRID.

Given the labels for all the cells (e.g., severe or mild COVID-19, dementia or not, or responding to immunotherapy or not), we first conducted differential expression analysis to identify differentially expressed genes. This step preselects the genes to facilitate model training since the original gene list is large. Specifically, we selected marker genes for each label/class separately. For example, in the case of severe and mild COVID-19, we performed a one-sided Mann-Whitney U rank test on each gene using the training data. To select the marker genes for severe (mild), the alternative hypothesis is that the expression in the severe (mild) group is stochastically greater than that in the other group. This test can also be generalized to a multi-class setting, with one group versus all other groups. We accounted for multiple testing by controlling the false discovery rate (FDR) using the Benjamini-Hochberg procedure and then selected the genes with FDR ≤ 0.05. As the number of marker genes may be large, we kept a maximum number of highly variable marker genes for each label, ranking the selected marker genes by normalized variance (computed using normalized data across all class labels). We selected at most 1,000 marker genes for each class. We then combined all the marker genes for different labels to create the gene list used later.

For the dimension reduction, we specified the number of components to be 40. For the unsupervised clustering using the Leiden algorithm to evaluate the latent components, we set the number of neighbors to 20 to compute a neighborhood graph and used a resolution value of 1.

The method was implemented using PyTorch. We used the Adam algorithm for optimization, with a learning rate of 0.0001 and weight decay of 0.0001. To avoid overfitting, we added dropout and early stopping. Dropout modules were added after each layer of the MLP, except the final layer, with a dropout probability of 0.5. We set the maximum number of epochs at 50,000 and the minimum number of epochs to 10,000. Training stops after no improvement in validation loss for 3,000 epochs.

When making predictions for each individual, we took the mean value of the scores across all their cells as the individual-level score.

### Implementation of sciRED and Spectra

We implemented sciRED following the workflow described in Pouyabahar et al. [9]. After selecting differentially expressed genes using the same procedure as in SDAN, we extracted the raw gene-level count data. Because sciRED explicitly models technical noise, no normalization or log transformation was applied prior to model fitting. For each gene, we fit a Poisson generalized linear model with an intercept, the cell’s library size, and protocol indicators (if available) as covariates. This procedure yielded estimated mean expression levels and Pearson residuals, where the residual matrix provides a variance-stabilized representation of gene expression that adjusts for sequencing depth and batch effects. The residual matrix was assembled into a dense array and standardized gene-wise to mean zero and unit variance. We then applied principal component analysis (PCA) to the residuals, retaining 40 components to match the number used for SDAN. To enhance interpretability, the PCA loadings were rotated using varimax, yielding approximately orthogonal gene programs. The rotated cell scores were then used as latent components for downstream cell-level classification, and individual-level scores were obtained by averaging cell-level predictions within each individual.

We implemented Spectra following the procedure described in [10]. If there were multiple cell types (e.g., CD4+ T cells and CD8+ T cells for COVID-19 severity study), we first constructed a union gene list by combining cell type–specific differentially expressed genes. Spectra was then trained on the merged dataset containing cells from all relevant cell types, using this union gene list for gene selection. The gene expression matrix was read-depth normalized and log-transformed, consistent with the preprocessing used for SDAN, and the same gene–gene adjacency matrix based on BioGRID interactions was used to encode gene annotation information. The Spectra model decomposes the expression matrix into two types of latent components: global (shared) factors and cell type-specific factors. In our studies, there were one or two cell types of interest. If there was only one cell type, we allocated 40 components, matching the number used in SDAN. If there were two cell types, we allocated 20 global components and 10 cell type-specific components for each cell type, following the structure suggested by Kunes et al. [10]. After fitting Spectra, we obtained a loading matrix mapping genes to latent components. We then projected the read-depth normalized and log-transformed gene expression data into the latent space using the learned loadings. These latent components were used for downstream cell-level classification, and individual-level scores were obtained by averaging cell-level predictions within each individual.

## Supporting information

Supplementary results

## Data Availability

### Su et al. COVID-19 dataset

Gene expression data were downloaded from ArrayExpress: https://www.ebi.ac.uk/arrayexpress/experiments/E-MTAB-9357/ on 2/22/2022. Patient information was extracted from Table S1 of Su et al. [14]: https://data.mendeley.com/datasets/tzydswhhb5/5 on 2/22/2022.

### SEA-AD dataset

The snRNA-seq data were downloaded from cellxgene websitehttps://cellxgene.cziscience.com/collections/1ca90a2d-2943-483d-b678-b809bf464c30 on 2021/11/21. Donor meta data and clinical data were downloaded from the SEA-AD website https://portal.brain-map.org/explore/seattle-alzheimers-disease/seattle-alzheimers-disease-brain-cell-atlas-download on 2022/1/7.

### Cancer immunotherapy dataset

The gene expression data of Sade-Feldman et al. 2018 [21] were downloaded from https://www.ncbi.nlm.nih.gov/geo/query/acc.cgi?acc=GSE120575. on 2023/11/6. Cell and sample information was extracted from Supplementary Table 1 of Sade-Feldman et al. 2018 [21]. The gene expression and meta-data of Yost et al. 2019 [22] were downloaded from https://www.ncbi.nlm.nih.gov/geo/query/acc.cgi?acc=GSE123813.

## Code Availability

Codes for data processing and running SDAN for all the datasets are available at https://github.com/Sun-lab/SDAN

## Key Points

- SDAN identifies gene sets that can classify cells and are composed of biologically related genes.
- Gene sets that classify cells are easier to interpret than a larger number of differentially expressed genes.
- In a deep learning model for genes, combining gene annotation and gene expression data is an effective solution to learn dependency between genes.

