## Supplementary results for "Supervised Deep Learning with Gene Annotation for Cell Classification"

Zhexiao Lin
Department of Statistics, University of California, Berkeley, CA

Yuanyuan Gao
Department of Statistics, University of California, Berkeley, CA

Wei Sun
Biostatistics Program, Fred Hutch Cancer Center, Seattle, WA
Department of Biostatistics, University of Washington, Seattle, WA
Department of Biostatistics, University of North Carolina, Chapel Hill, NC

January 13, 2026

#### 1 Prediction of COVID-19 severeness

##### 1.1 Data processing

Su et al. (2020) collected scRNA-seq data of around 0.55 million PBMCs from 254 COVID-19 samples at two time points: baseline (BL, initial clinical diagnosis) and AC (a few days after initial clinical diagnosis). Our objective was to predict COVID-19 severity using 129 baseline samples. We identified CD4+ T and CD8+ T cells using paired single-cell TCR data. The gene expression data, downloaded from ArrayExpress, were already read-depth corrected and log-transformed. We converted the expression of gene $j$ in cell $i$, denoted by $x_{ij}$, to count data using the following transformation: $y_{ij} = (\exp(x_{ij}) - 1)/d_i$ where $d_i = \exp(\min_j(x_{ij})) - 1$ is the read-depth. After removing a small proportion of cells with relatively high expression from mitochondrial genes, there were 73,383 CD4+ T cells and 40,223 CD8+ T cells. Both CD4+ T and CD8+ T cells contain 24,966 genes. We normalized each cell such that the total count of cells was 10,000, and log-transformed the data. We excluded all the mitochondrial genes in our downstream analysis and only kept genes expressed in more than 2% of cells. Then the number of genes remaining was 7,704 for CD4+ T cells and 7,864 for CD8+ T cells. To construct the label of disease severity, we used the World Health Organization [WHO] ordinal scale [WOS] = 1-2 for mild, WOS = 3–4 for moderate, and WOS = 5–7 for severe. Each cell was labeled based on the corresponding patient's label. We only used the data from 49 mild and 32 severe patients for our analysis. There were 25,687 and 16,334 CD4+ T cells from mild and severe patients, respectively; and 16,449 and 7,671 CD8+ T cells from mild and severe patients, respectively. We randomly split the patients in the severe group and the mild group into two subsets of equal sizes and used the data from one subset of patients as training data and the remaining data as testing data. For this and all the other datasets, we randomly took 10% of the cells in the training data as the validation data and used the remaining cells for training.

#### 1.2 Additional results

For CD8+ T cells, we found 649 differentially expressed genes (FDR $\leq 0.05$) with higher expression in mild cases and 2,926 differentially expressed genes with higher expression in severe cases. By default, we kept at most 1,000 marker genes with the highest variation for each class, which led to 1,649 genes (649 for mild and 1,000 for severe) in total. Varying the weight parameter in the set of {0, 0.25, 0.5, 1, 2, 5, 10}, the AUC scores at the cell level were {0.898, 0.890, 0.879, 0.884, 0.878, 0.857, 0.851}. At the individual level, the AUC scores were {0.967, 0.970, 0.976, 0.958, 0.964, 0.973, 0.955}.

We used the goseq function in R (Young et al., 2010) to assess whether each gene set identified by SDAN was enriched with genes from some functional categories. We considered three types of functional categories downloaded from MSigDB (Liberzon et al., 2015):

- Gene ontology terms (biological processes): `c5.go.bp.v2023.2.Hs.symbols.gmt`.
- Reactome pathways: `c2.cp.reactome.v2023.2.Hs.symbols.gmt`.
- Immune-related categories: `c7.all.v2023.2.Hs.symbols.gmt`.

Multiple tests across functional categories were accounted for by calculating the Benjamini & Hochberg False Discovery Rate (FDR).

The number of enriched function categories increases as the weight on the unsupervised loss function increases from 0.25 to 2 (Supplementary Figure 1). There is no enriched functional category when the weight is 0. This is expected because a larger weight on the unsupervised loss encourages the selection of gene sets that have more annotated gene-gene interactions within each gene set.

For CD4+ T cells, we identified 1,126 differentially expressed genes for mild cases and 2,057 for severe cases. From these, we selected 2,000 genes, with 1,000 for each class. We varied the weight parameter across the values {0, 0.25, 0.5, 1, 2, 5, 10} and observed the following AUC scores at the cell level: {0.907, 0.914, 0.909, 0.906, 0.897, 0.884, 0.829}. At the individual level, the AUC scores were {0.940, 0.940, 0.952, 0.958, 0.949, 0.952, 0.955}.

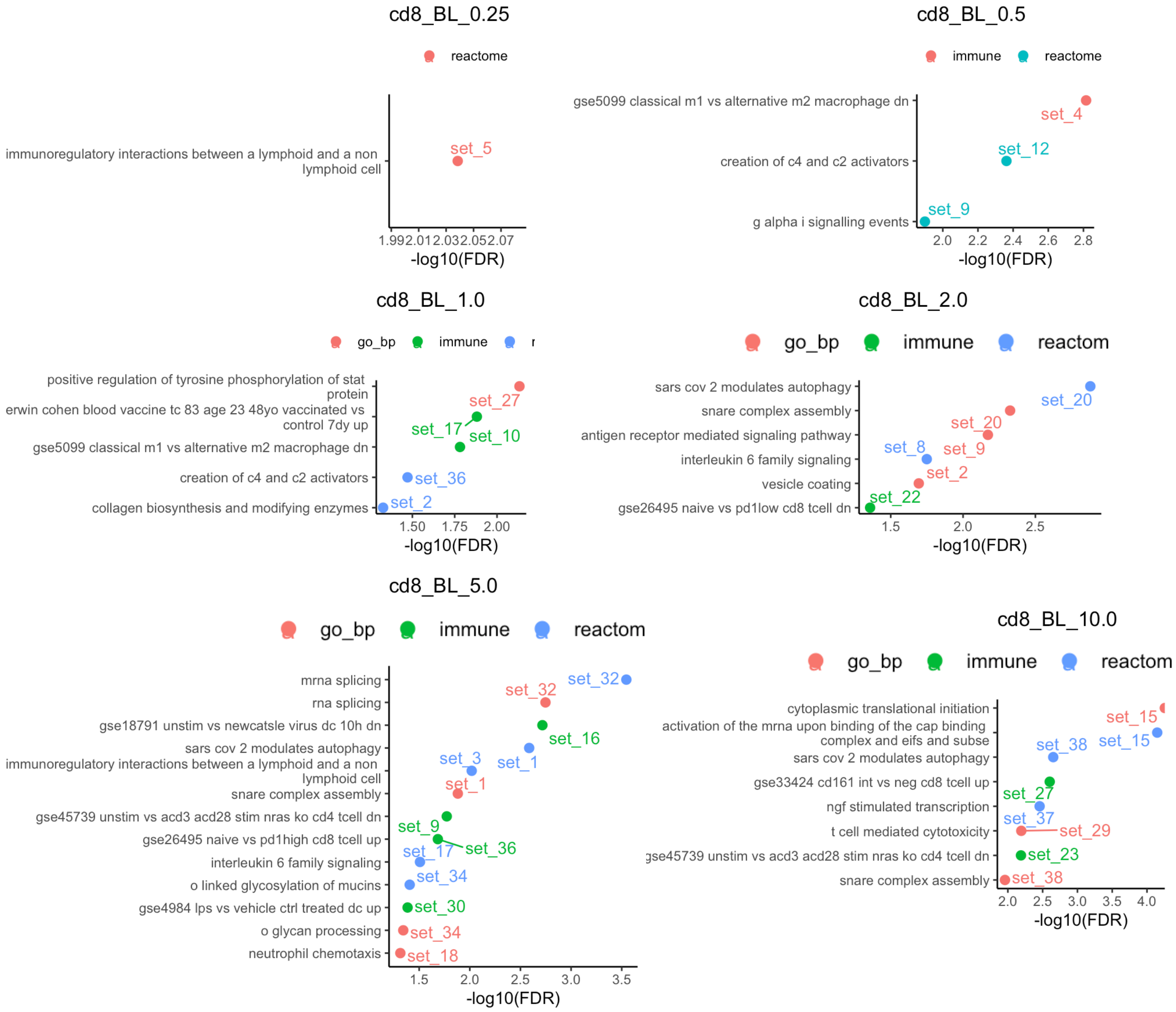


Supplementary Figure 1: Functional category enrichment analysis results using gene sets identified by CD8+ T cells with different weights on the unsupervised loss.

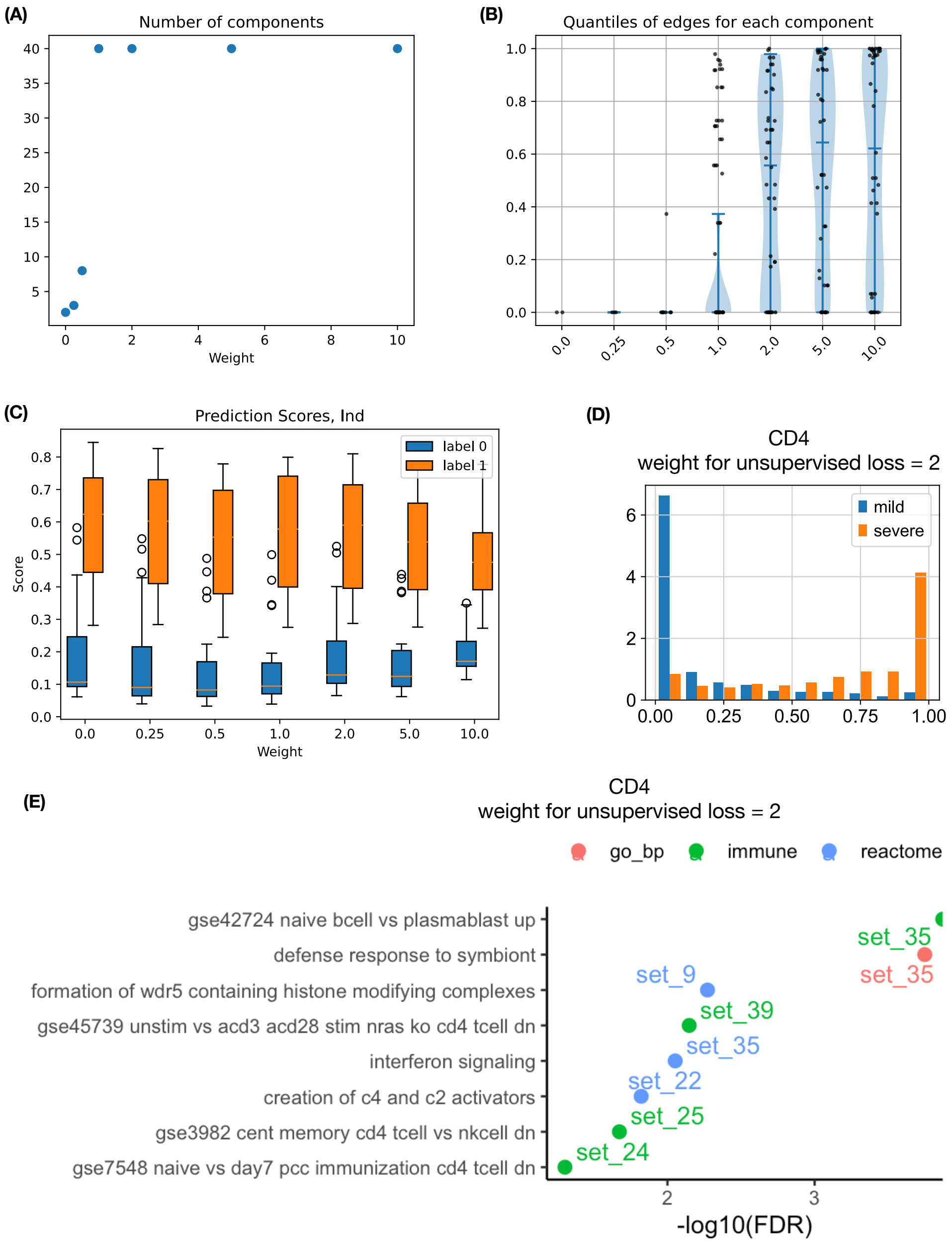


Supplementary Figure 2: Summary of results for CD4+ T cells. **(A)** The number of latent components for different weights for the unsupervised loss. **(B)** Violin plots of connection quantiles. Each point is one component, the the connection quantile is computed based on the number of connections from randomly selected gene sets. **(C)** Boxplots of **individual**-level scores (average of cell level scores) for severe disease, across different weights for the unsupervised loss. **(D)** Distribution of cell-level prediction scores (for CD8+ T cells using weight 2.0) for severe disease in testing data, stratified by the status of severe/mild diseases of the corresponding individual. **(E)** Enriched function categorises of identified gene sets when using weight 2 for the unsupervised loss.

### 2 Prediction of dementia

We used the single nucleotide RNA-seq (snRNA-seq) data from Seattle Alzheimer's Disease Cell Atlas (SEA-AD) consortium (Gabitto et al., 2023). This dataset consisted of multi-omic data from middle temporal gyrus (MTG) of 84 donors. MTG is a brain area that is relevant to dementia since it is involved in language and semantic memory processing and higher order visual processing. We studied gene expression of two cell types: astrocyte (Astro) and microglia (Micro-PVM). There were 70,009 Astro cells and $\sim$ 40,000 Micro-PVM cells. Both Astro and Micro-PVM data contained 36,517 genes. We normalized the gene expression of each cell such that the total UMI count of each cell is 10,000, and then log-transformed the data. We excluded all mitochondria genes and only kept genes expressed in more than 2% of the cells. The number of genes remained is 16,058 for Astro and 13,895 for Micro-PVM. We labeled each cell based on the cognitive status of the donor. There were 42 donors with dementia and 42 without dementia. We randomly selected half of dementia donors and non-dementia donors to construct the training data and used the remaining donors' data as testing data.

For Astro cells, we identified 6,765 differentially expressed genes for non-dementia class and 2,120 for dementia class. From these, we selected 2,000 genes. We varied the weight parameter across the values $\{0, 0.25, 0.5, 1, 2, 5, 10\}$ and observed the following AUC scores at the cell level: $\{0.740, 0.726, 0.726, 0.722, 0.715, 0.677, 0.667\}$. At the individual level, the AUC scores were $\{0.748, 0.739, 0.748, 0.751, 0.744, 0.744, 0.746\}$ (Supplementary Figure 3).

For Micro-PVM cells, we found 2,554 differentially expressed genes for non-dementia class and 1,939 for dementia class. Fropm them, we selected 2,000 genes. Varying the weight parameter across the same range, the AUC scores at the cell level were $\{0.648, 0.650, 0.643, 0.650, 0.634, 0.620, 0.602\}$. At the individual level, the AUC scores were $\{0.719, 0.714, 0.726, 0.737, 0.735, 0.728, 0.730\}$ (Supplementary Figure 4).

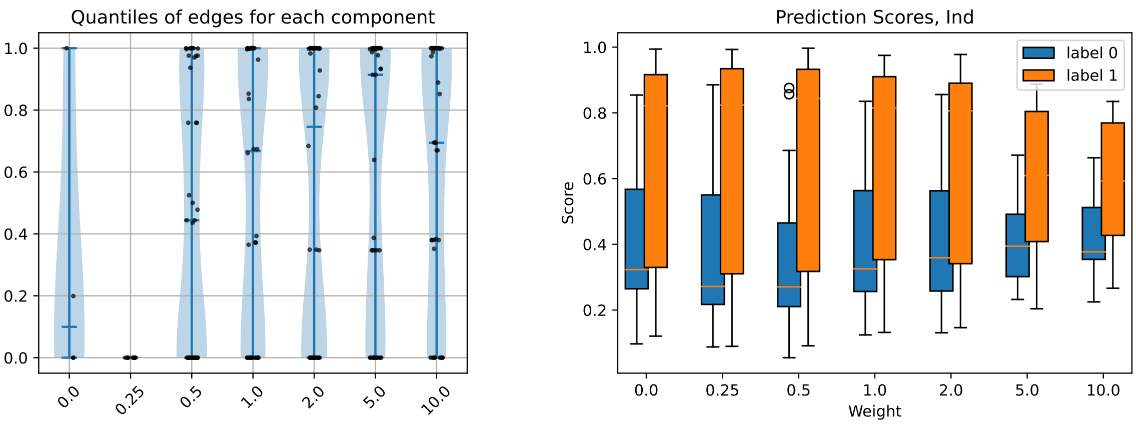


(A) (B)

Supplementary Figure 3: Exploration of different weights for unsupervised loss using snRNA-seq data from **Astrocyte**. (A) Violin plots of connection quantiles. Each point is one component. The connection quantile (y-axis) is computed based on the number of connections from randomly selected gene sets. (B) Boxplots of individual-level SDAN prediction scores (average of cell level scores) across different weights for the unsupervised loss.

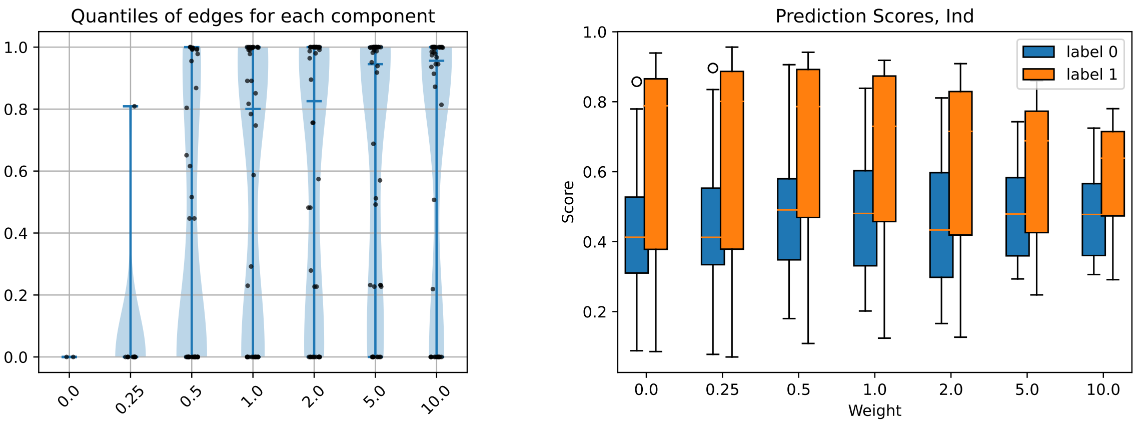


(A) (B)

Supplementary Figure 4: Exploration of different weights for unsupervised loss using snRNA-seq data from **Microglia-PVM**. (A) Violin plots of connection quantiles. (B) Boxplots of individual-level SDAN prediction scores.

### 3 Prediction of patient response to cancer immunotherapy

Due to the limited sample size of existing datasets, we combined two datasets to predict cancer immunotherapy response. The training data were from Sade-Feldman et al. (2018), which consisted of 6,350 CD8+ T cells and 36,602 genes from 17 responding tumors and 31 non-responding tumors. The data were already normalized and thus we directly used them. We excluded all mitochondria genes and kept genes expressed in more than 2% cells. The number of genes left was 13,480. We used the corresponding patients' response to immunotherapy to label all the cells. All the data from Sade-Feldman et al. (2018) were used for training. We tested the performance using Yost et al. (2019) data, which consisted of 27,924 CD8+ T cells and 18,189 genes from 15 patients, with 8 responders and 7 non-responders.

We identified 7,112 differentially expressed genes for no-response cases and 259 for response cases. From these, we selected 1,259 genes. We varied the weight parameter across the values {0, 0.25, 0.5, 1, 2, 5, 10} and observed the following AUC scores at the cell level: {0.572, 0.587, 0.590, 0.518, 0.532, 0.531, 0.553}. At the individual level, the AUC scores were {0.732, 0.821, 0.804, 0.643, 0.661, 0.482, 0.607} (Supplementary Figure 5).

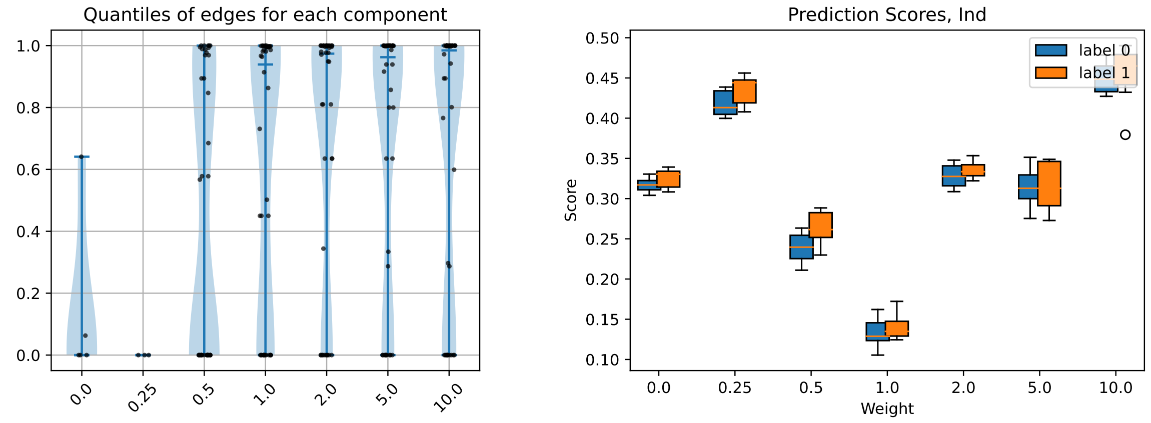


(A) (B)

Supplementary Figure 5: Exploration of different weights for unsupervised loss using scRNA-seq data of CD8+ T cells to predict cancer immunotherapy response . (A) Violin plots of connection quantiles. (B) Boxplots of individual-level SDAN prediction scores.

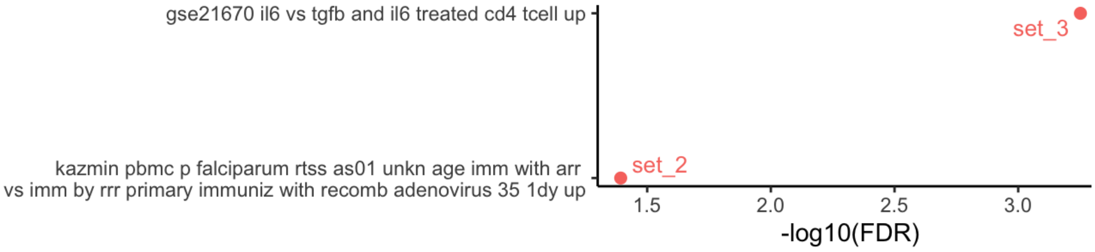


(A) Weight 0.25 for unsupervised loss.

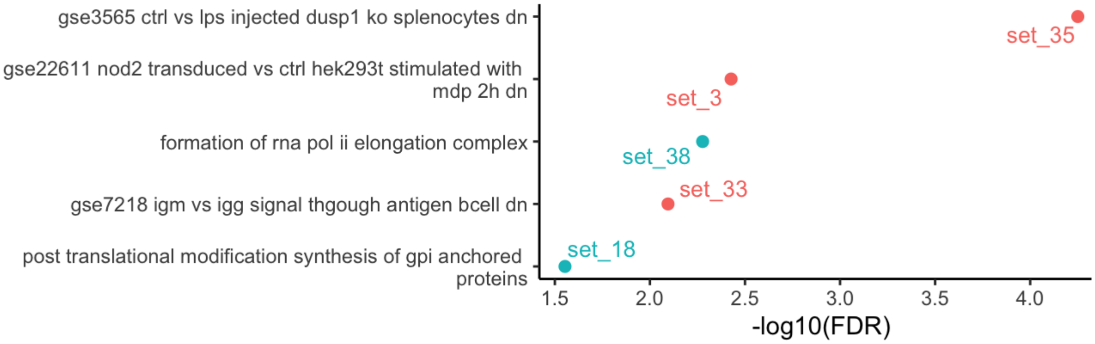


(B) Weight 0.5 for unsupervised loss.

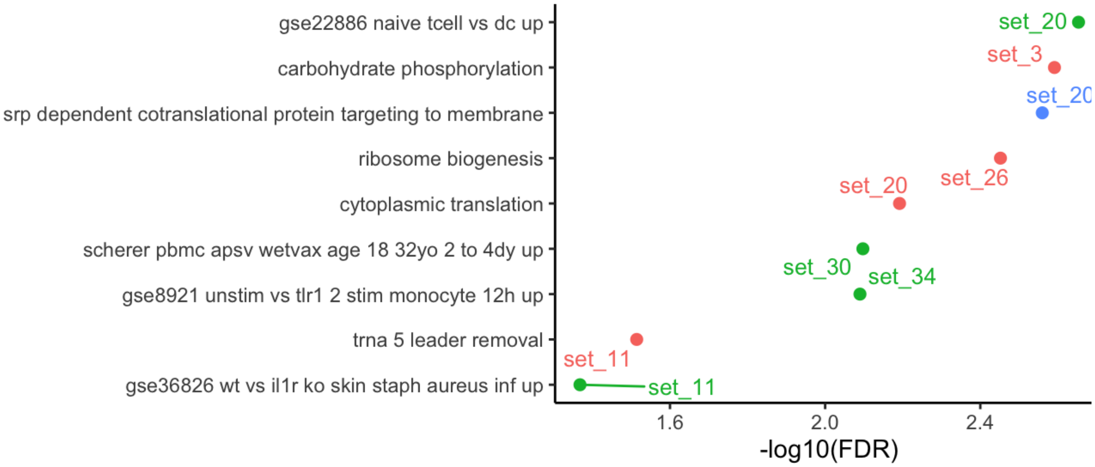


(C) Weight 1.0 for unsupervised loss.

Supplementary Figure 6: Functional category enrichment analysis across different weights for unsupervised loss using scRNA-seq data of CD8+ T cells to predict cancer immunotherapy response.

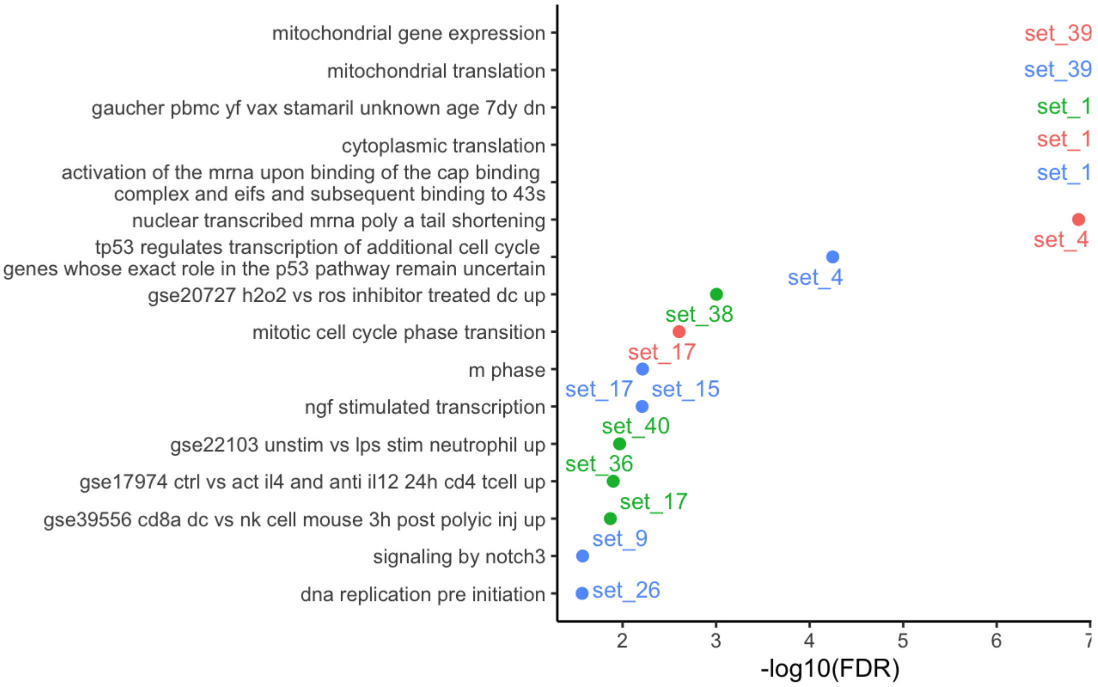


(A) Weight 5.0 for unsupervised loss.

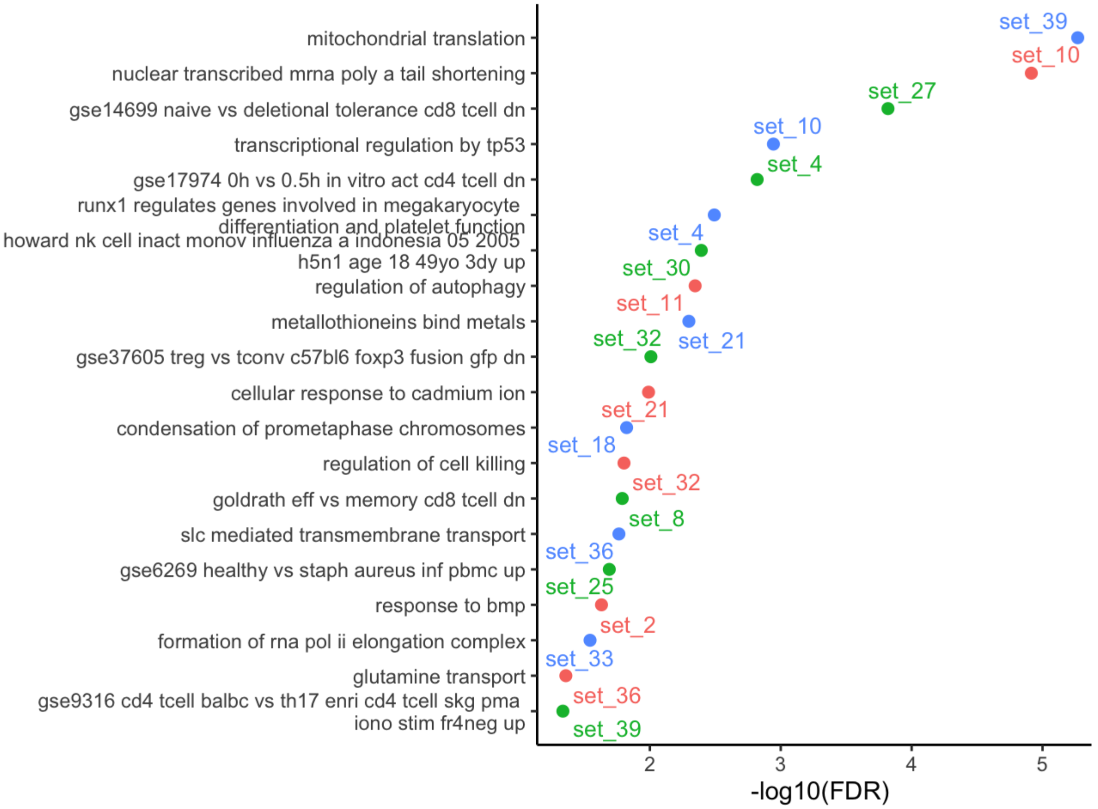


(B) Weight 10.0 for unsupervised loss.

Supplementary Figure 7: Continuation of functional category enrichment analysis across different weights for unsupervised loss using scRNA-seq data of CD8+ T cells to predict cancer immunotherapy response.

| hgnc_symbol | chr | description |
|---|---|---|
| ATPAF1 | chr1 | ATP synthase mitochondrial F1 complex assembly factor 1 |
| CDK5RAP1 | chr20 | CDK5 regulatory subunit associated protein 1 |
| DHX58 | chr17 | DExH-box helicase 58 |
| EARS2 | chr16 | glutamyl-tRNA synthetase 2, mitochondrial |
| GTPBP10 | chr7 | GTP binding protein 10 |
| KHSRP | chr19 | KH-type splicing regulatory protein |
| MALSU1 | chr7 | mitochondrial assembly of ribosomal large subunit 1 |
| MCUB | chr4 | mitochondrial calcium uniporter dominant negative subunit beta |
| MRPL23 | chr11 | mitochondrial ribosomal protein L23 |
| MRPL48 | chr11 | mitochondrial ribosomal protein L48 |
| MRPL55 | chr1 | mitochondrial ribosomal protein L55 |
| MRPS33 | chr7 | mitochondrial ribosomal protein S33 |
| MRRF | chr9 | mitochondrial ribosome recycling factor |
| MTFMT | chr15 | mitochondrial methionyl-tRNA formyltransferase |
| MTIF2 | chr2 | mitochondrial translational initiation factor 2 |
| MTO1 | chr6 | mitochondrial tRNA translation optimization 1 |
| MTRES1 | chr6 | mitochondrial transcription rescue factor 1 |
| NDUFAF4 | chr6 | NADH:ubiquinone oxidoreductase complex assembly factor 4 |
| NDUFAF5 | chr20 | NADH:ubiquinone oxidoreductase complex assembly factor 5 |
| NDUFS4 | chr5 | NADH:ubiquinone oxidoreductase subunit S4 |
| PHAX | chr5 | phosphorylated adaptor for RNA export |
| SARS2 | chr19 | seryl-tRNA synthetase 2, mitochondrial |
| ZNF431 | chr19 | zinc finger protein 431 |

Supplementary Table 1: Gene set 1 identified by SDAN.

### 4 Comparisons with other methods

#### 4.1 Metrics

To quantify the sparsity of the learned loading vectors for each gene, we use *Hoyer sparsity*, a scale-invariant measure that captures the extent to which a vector's mass is concentrated on a small number of entries. For a vector $\mathbf{v} \in \mathbb{R}^d$, Hoyer sparsity is defined in terms of its $L_1$ and $L_2$ norms as

$$Sparsity(\mathbf{v}) = \frac{\sqrt{d} - \frac{\|\mathbf{v}\|_1}{\|\mathbf{v}\|_2}}{\sqrt{d} - 1},$$

which takes values in $[0, 1]$. Values close to 0 correspond to dense vectors with relatively uniform weights, whereas values close to 1 indicate highly sparse vectors dominated by a small number of nonzero entries. Importantly, Hoyer sparsity is invariant to scaling of the vector, making it suitable for comparing sparsity across methods with different normalization schemes. In our analysis, we compute Hoyer sparsity for each gene based on its loading vector across latent components in the learned loading matrix. Higher Hoyer sparsity thus reflects more interpretable representations, as the gene loads on only a small subset or even a single latent factor.

#### 4.2 Results

Supplementary Table 2: Summary of Logistic Regression AUC for COVID-19 severity prediction (Su et al., 2020) using GNN Across Weights. The setting with optimal performance is labeled by red color.

| **Weight** | **CD4 Cell** | **CD8 Cell** | **CD4 Ind** | **CD8 Ind** |
|---|---|---|---|---|
| 0.00 | 0.9070 | 0.8983 | 0.9405 | 0.9673 |
| 0.25 | 0.9138 | 0.8904 | 0.9405 | 0.9702 |
| 0.50 | 0.9095 | 0.8791 | 0.9524 | 0.9762 |
| 1.00 | 0.9063 | 0.8838 | 0.9583 | 0.9583 |
| 2.00 | 0.8967 | 0.8785 | 0.9494 | 0.9643 |
| 5.00 | 0.8839 | 0.8566 | 0.9524 | 0.9732 |
| 10.00 | 0.8290 | 0.8506 | 0.9554 | 0.9554 |

Supplementary Table 3: Summary of Logistic Regression AUC for Dementia study (Gabitto et al., 2023) using GNN Across Weights. The setting with optimal performance is labeled by red color.

| Weight | Astro Cell | Micro Cell | Astro Ind. | Micro Ind. |
|---|---|---|---|---|
| 0.00 | 0.7397 | 0.6479 | 0.7483 | 0.7188 |
| 0.25 | 0.7260 | 0.6499 | 0.7392 | 0.7143 |
| 0.50 | 0.7257 | 0.6431 | 0.7483 | 0.7256 |
| 1.00 | 0.7216 | 0.6505 | 0.7506 | 0.7370 |
| 2.00 | 0.7151 | 0.6342 | 0.7438 | 0.7347 |
| 5.00 | 0.6766 | 0.6204 | 0.7438 | 0.7279 |
| 10.00 | 0.6672 | 0.6021 | 0.7460 | 0.7302 |

Supplementary Table 4: Summary of Logistic Regression AUC for prediction ICI response (Yost et al., 2019) using GNN Across Weights. The setting with optimal performance is labeled by red color.

| Weight | CD8T Cell | CD8T Ind. |
|---|---|---|
| 0.00 | 0.5720 | 0.7321 |
| 0.25 | 0.5867 | 0.8214 |
| 0.50 | 0.5900 | 0.8036 |
| 1.00 | 0.5180 | 0.6429 |
| 2.00 | 0.5316 | 0.6607 |
| 5.00 | 0.5310 | 0.4821 |
| 10.00 | 0.5527 | 0.6071 |

Supplementary Table 5: Comparison of Logistic Regression AUC at Cell and Individual Levels (GNN Weight = 2.0). The setting with optimal performance is labeled by red color.

| Method | Level | CD4+ T | CD8+ T | Astro | Micro-PVM | CD8+ T |
|---|---|---|---|---|---|---|
| GNN | Cell | 0.8967 | 0.8785 | 0.7151 | 0.6342 | 0.5501 |
| | Individual | 0.9494 | 0.9643 | 0.7438 | 0.7347 | 0.6429 |
| Spectra | Cell | 0.8039 | 0.7696 | 0.7788 | 0.6832 | 0.5424 |
| | Individual | 0.9315 | 0.9256 | 0.7324 | 0.7664 | 0.5357 |
| sciRED | Cell | 0.8902 | 0.8839 | 0.7505 | 0.6720 | 0.5141 |
| | Individual | 0.9554 | 0.9554 | 0.7483 | 0.7574 | 0.6429 |

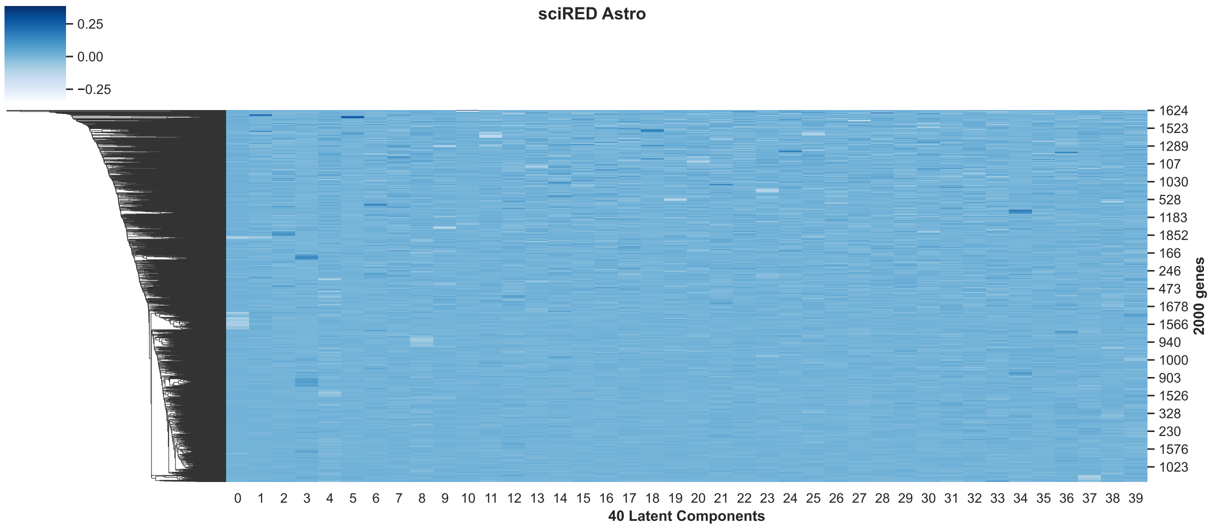


Supplementary Figure 8: Gene assignment scores to different components identified by sciRED using gene expression data of astrocyte (Gabitto et al., 2023).

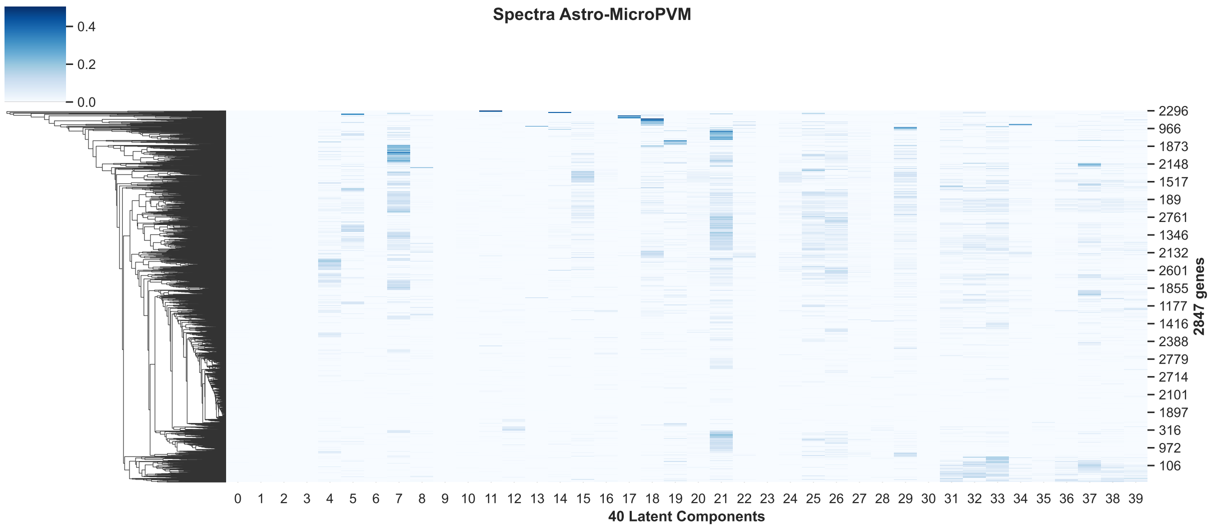


Supplementary Figure 9: Gene assignment scores to different components identified by Spectra using gene expression data of astrocyte.

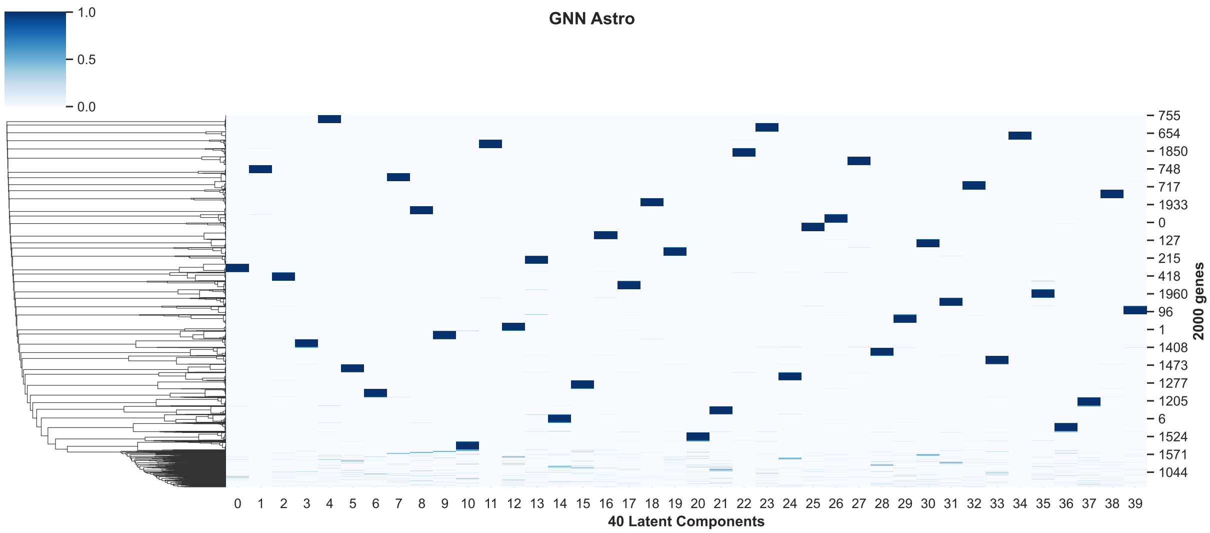


Supplementary Figure 10: Gene assignment scores to different components identified by SDAN using gene expression data of astrocyte.
